# The role of Kabuki Syndrome genes *KMT2D* and *KDM6A* in development: Analysis in Human sequencing data and compared to mice and zebrafish

**DOI:** 10.1101/2020.04.03.024646

**Authors:** Rwik Sen, Ezra Lencer, Elizabeth A. Geiger, Kenneth Jones, Tamim H. Shaikh, Kristin Bruk Artinger

## Abstract

*KMT2D* and *KDM6A* are epigenetic regulators that have been implicated in Kabuki Syndrome, a rare congenital birth defect with multiple tissue and organ abnormalities, including craniofacial and heart defects. Our previous study identified human families with mutations in the epigenetic modifiers KMT2D and KDM6A, which is implicated in 32% and 10% of Kabuki Syndrome patients respectively. To understand the connection to Kabuki syndrome patients, and the transcriptional targets of KMT2D and KDM6A in humans, we performed RNA sequencing (seq) of lymphoblastoid cells from Kabuki Syndrome patients carrying mutations in *KMT2D* and *KDM6A*. We identified 1995 significant changes in transcriptional targets for *KMT2D* and 1917 for *KDM6A*, as compared to control. When compared with RNA-seq datasets obtained from other mouse and zebrafish studies, our analysis revealed *KMT2D* mutations affect the expression of 76 orthologous genes across all three datasets. Similarliy, *KDM6A* afftects the expprssion of 7 orthologous genes across three datasets. Despite the differences in cell types, stages, and species in the comparison between the transcriptomic datasets, there are common gene expression changes associated with KMT2D and KDM6A mutations. qPCR on novel zebrafish mutants confirmed the differentially expression in KMT2D or KDM6A mutant backgrounds. Taken together, our results show that *KMT2D* and *KDM6A* regulate common and unique genes across humans, mice, and zebrafish for early craniofacial and cardiac development and this information contributes to the understanding of epigenetic dysregulation during development of Kabuki syndrome.

## Introduction

Kabuki syndrome is a rare congenital abnormality with characteristic facial features superficially similar to the makeup used by actors in traditional Japanese theatre where the name arises. The craniofacial malformation results from defects in the eyelid tears, arched and broad eyebrows, depressed nasal tip, and prominent or cupped ears (Adam et al., 2019). These patients also present with cardiac issues as well as growth defects such as short stature and microcephy. Approximately 28-80 % of Kabuki Syndrome patients are afflicted with congenital heart defects (Digilio et al., 2017). Common heart defects include atrial and ventricular septal defects, bicuspid aortic valves, aortic coarctation, double outlet right ventricle, transposition of great arteries, infundibular pulmonary stenosis, dysplastic mitral valve, Tetralogy of Fallot, etc. (Cocciadiferro et al., 2018; Digilio et al., 2017; Shangguan et al., 2019; Yap et al., 2020).

The combination of defects observed in Kabuki Syndrome are similar to those observed in embryos with deficiencies in neural crest cells (NCCs), a multipotent population of cells that give rise to craniofacial cartilage and bone, cardiac outflow track as well as multiple other derivatives. NCCs form early in development as part of the central nervous system, and then migrate long distances to begin a differentiation program. Because they undergo multiple morphogenetic and cellular processes, NCCs rely on a precise gene and epigenetic regulatory network for their development. Several epigenetic processes are important for development including DNA methylation (Serra-Juhe et al., 2015), histone modifications (Zaidi et al., 2013; Q. J. Zhang & Liu, 2015), chromatin remodeling (Bruneau, 2010; Delgado-Olguin, Takeuchi, & Bruneau, 2006), and microRNA (Toni, Hailu, & Sucharov, 2020) and contribute to the etiology of developmental disorders. Because of the ability to transcriptionally regulate many genes at the same time, defects in epigenetic regulators help to explain why patients with the same genotype and syndrome often have large variations in phenotype, penetrance, and severity. Indeed, mutations in chromatin remodeler Chd7 and histone acetyltransferase Kat6a/b are associated with birth defects leading to Charge and Ohdo syndromes, respectively (Vissers et al., 2004) (Campeau et al., 2012).

*KMT2D*, also known as MLL2 methylates histone 3 at lysine 4 (H3K4) which is associated with transcriptional repression, and *KDM6A*, demethylates histone 3 at lysine 27 (H3K27), mostly associated with transcriptional actiation (Ali, Hom, Blakeslee, Ikenouye, & Kutateladze, 2014; Hong et al., 2007; Koutsioumpa et al., 2019; Lan et al., 2007), both genes are frequently mutated in patients with Kabuki Syndrome (Cocciadiferro et al., 2018; Gazova, Lengeling, & Summers, 2019; Luperchio, Applegate, Bodamer, & Bjornsson, 2019; Shangguan et al., 2019; Tekendo-Ngongang, Kruszka, Martinez, & Muenke, 2019; Yap et al., 2020) (Ang et al., 2016; Lee, Lee, & Lee, 2012; Schwenty-Lara, Nurnberger, & Borchers, 2019; Serrano, Demarest, Tone-Pah-Hote, Tristani-Firouzi, & Yost, 2019). *KMT2D* or lysine-specific methyltransferase 2D, is also known as *MLL2* or myeloid/lymphoid or mixed-lineage leukemia 2. It belongs to a family of 7 SET1-like histone methyltransferases (Ali et al., 2014). KMT2D contains a SET domain for histone methylation, PHD and coiled-coil domains, and zinc finger domains (Ali et al., 2014). The PHD domain of zebrafish *kmt2d*, which shares 86.1% identity with human *KMT2D (***Figure 1A**).

**Figure 1:**
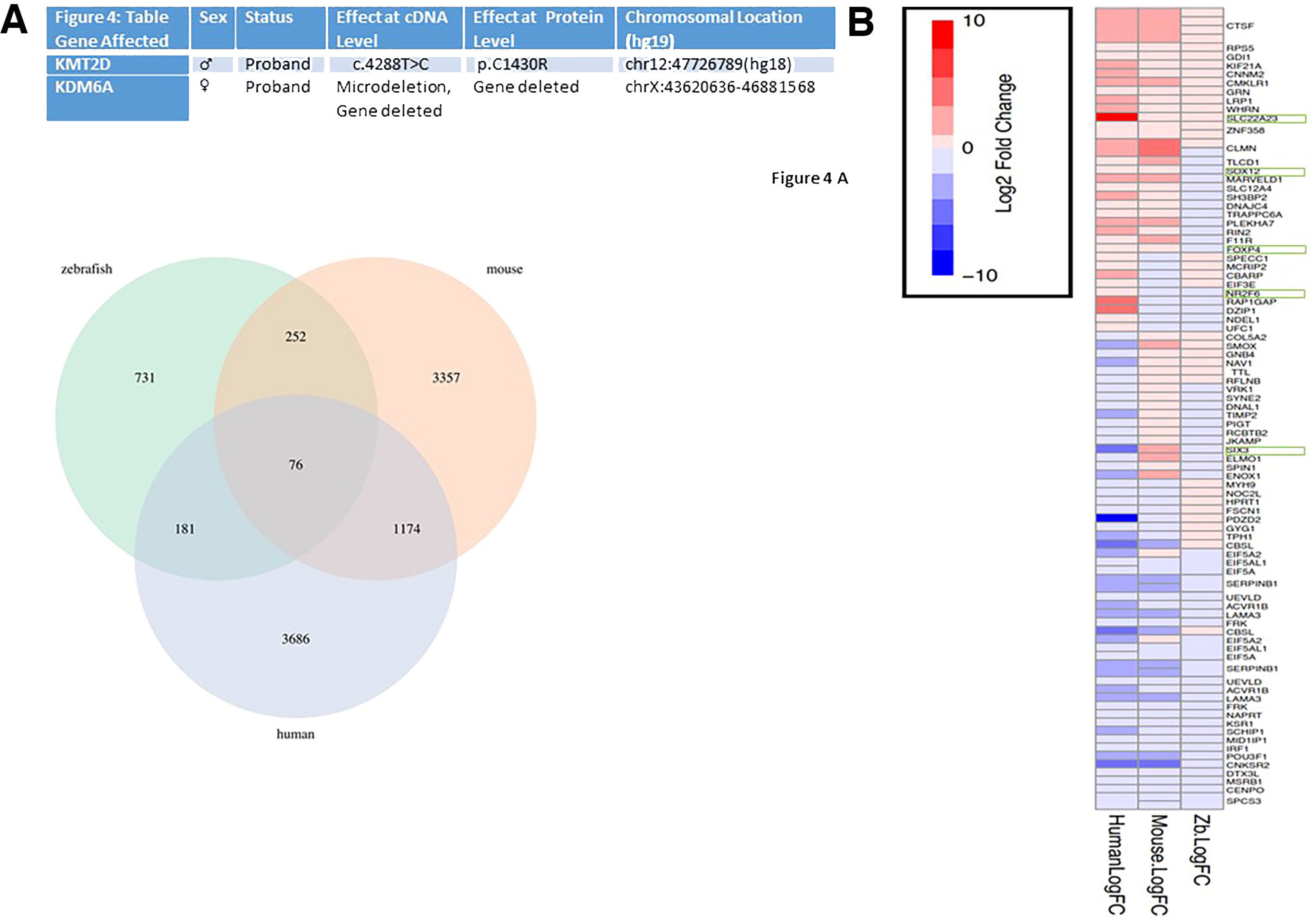
*KMT2D* shares common targets across humans, mice and zebrafish. Table: Mutation information of patients. A. Venn diagrams showing overlap between gene lists derived from RNA-sequencing (seq) analysis of *KMT2D* mutant datasets from humans, mice, and zebrafish as labeled. B. Heat maps showing 76 common targets among human, mice, and zebrafish. Genes selected for RT-qPCR in Figure 1 C. are highlighted in green boxes.

**Figure 2:**
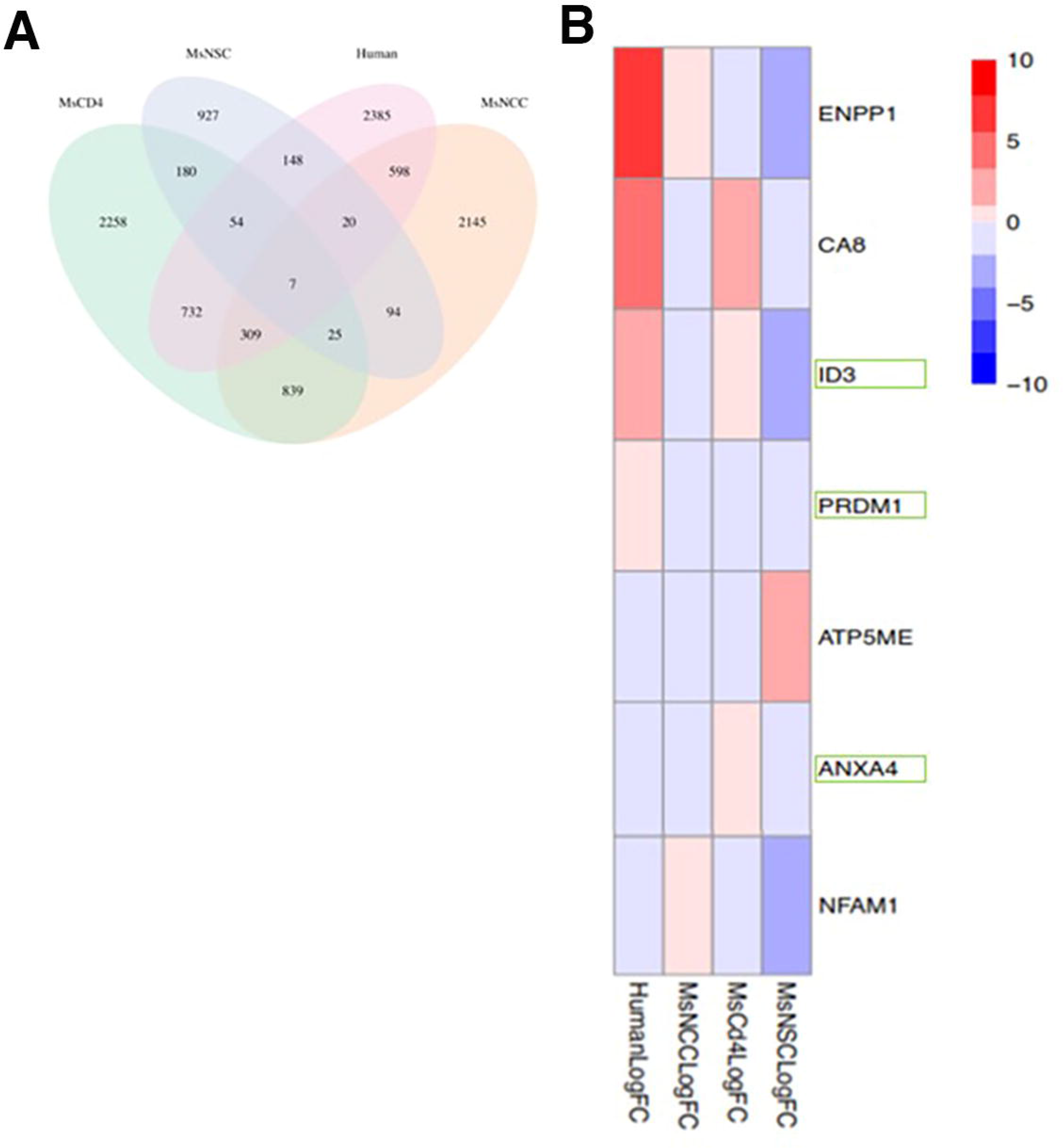
*KDM6A* shares common targets across humans and various cell types of mice, which are similarly regulated in zebrafish. A. Venn diagrams showing overlap between gene lists derived from RNA-seq analysis of *KDM6A* mutant datasets from humans and three distinct cell types of mice, as labeled. B. Heat maps showing 7 common targets among human and 3 cell types of mice. Abbreviations: Ms – mouse, NCC – neural crest cells, NSC – neural stem cells, Cd4 – Cd4 T lymphocytes. Genes selected for RT-qPCR in Figure 2 C. are highlighted in green boxes.

Recent studies have significantly contributed to our understanding of cardiac development and function as regulated by *KMT2D* (Ang et al., 2016; Schwenty-Lara, Nurnberger, et al., 2019; Serrano et al., 2019), and *KDM6A* (Lee et al., 2012). The PHD domain of zebrafish *kmt2d*, which shares 86.1% identity with human *KMT2D*, was targeted by CRISPR-mediated gene disruption in a recent study on *kmt2d* in heart development (Serrano et al., 2019). The study generated 3 alleles with premature stop codons that cause a truncation of the PHD domain (Serrano et al., 2019) and categorized phenotypes, including heart edema and microcephaly, and a shorter body axis (Serrano et al., 2019). They report a robust Notch signaling hyperactivation in endocardial and endothelial cells that affect vasculogenesis and angiogenesis, and a rescue of cardiovascular phenotype by pharmacological inhibition of Notch signaling (Serrano et al., 2019). Another recent study on *kmt2d* showed morpholino-based knockdown of *kmt2d* in *Xenopus* causes heart defects (Schwenty-Lara, Nurnberger, et al., 2019), and impairs NCC development (Schwenty-Lara, Nehl, & Borchers, 2019). In mouse studies, development of limbs, palate, central nervous system, and heart are impaired upon mutating *Kmt2d*, with hypoplasia in frontonasal bone, fully penetrant cleft palate, mandible hypoplasia, deficits in palatal shelf elevation and cranial base ossification (Fahrner et al., 2019; Shpargel, Mangini, Xie, Ge, & Magnuson, 2020). Further, disruption of *Kmt2d* show disorganized interventricular septum, hypoplasia of compact myocardium indicating decreased cardiomyocyte proliferation and failure of outflow tract septation, suggesting a role of *Kmt2d* in cardiac development (Ang et al., 2016).

*KDM6A* (and *kdm6l* in zebrafish) or Lysine Demethylase 6A, is also known as *UTX* or Ubiquitously transcribed tetratricopeptide repeat, X chromosome. UTX, UTY and JMJD3 comprise a subfamily of proteins containing JmjC-domain (Jumonji C) and zinc fingers, and they are evolutionarily conserved from *Caenorhabditis elegans* to human (Klose, Kallin, & Zhang, 2006; Lan et al., 2007; Shi & Whetstine, 2007). The JmjC-domain is associated with histone demethylation (Accari & Fisher, 2015), while the tetratricopeptide repeats at N-terminal regions of UTX and UTY are predicted protein interaction motifs (Lan et al., 2007). It is important to note that *KDM6A* or *UTX* is an X-linked gene, with 2 copies in females (XX) and 1 copy in males (XY), where the Y chromosome has the homolog, *UTY* (Itoh et al., 2019). Studies indicate that *UTY* and *UTX* diverged from a common ancestor, yet the functional role of *UTY* in histone demethylation needs further investigation, because some studies show that it has a lower histone demethylation activity than *UTX* (Faralli et al., 2016; Hong et al., 2007; Itoh et al., 2019; Lan et al., 2007; Shpargel, Starmer, Wang, Ge, & Magnuson, 2017; Shpargel, Starmer, Yee, Pohlers, & Magnuson, 2014; Walport et al., 2014).

To determine the effect of patient mutations on the transcriptome, we performed RNA-seq analysis on human patient lymphoblastoid cells (LBCs). We determined that 1995 genes were differentially regulated for *KMT2D*, and 1917 for *KDM6A*, as compared to control LBCs. We then compared common gene targets of *KMT2D* and *KDM6A* from published zebrafish, mice, with our human RNA-seq datasets in order to prioritize downstream genes for further study. Our results reveal that some of the genes which are involved in cardiogenesis, cardiac function, vasculature, neurogenesis, and overall development, are regulated by *KMT2D* and *KDM6A* across zebrafish, mice, and humans. Our *in vivo* analysis in CRISPR-mutated zebrafish indicate that *kmt2d* and *kdm6a* regulate genes involved in early embryonic, cardiac, and NCC development. Our results support the idea that mutations in zebrafish KMT2D and KDM6A have defects similar to Kabuki Syndrome patients, supporting previous studies in the field, identifies targets in human patients LBCs, and presents a pan-species perspective of the roles of *KMT2D* and *KDM6A* in development.

## Materials and Methods

### Human Cell Samples

Patient-derived lymphoblastoid cell lines (LCLs) used in this study were previously generated from KS patients with mutations in *KMT2D*/*MLL2* and *KDM6A* (Van Laarhoven et al 2015). Control LCLs from age and demography-matched subjects, were obtained from the Coriell Institute for Medical Research https://www.coriell.org/

### RNAseq of human Kabuki Syndrome patients and comparison across taxa

Human LCLs from *KMT2D* and *KDM6A* patients and controls, as mentioned above, were grown to a count of 10^6^ and pelleted. RNA was extracted by Zymo Research Direct-zol^™^ RNA MiniPrep kit (Catalog #R2050). Two cDNA libraries for each genotype (e.g. wildtype, *KMT2D* mutant, *KDM6A* mutant) were prepared from separate RNA MiniPreps using the Nugen mRNA kit for Illumina sequencing. Sequencing was performed on an Illumina NovaSEQ6000 (150bp paired-end reads) at the Genomics and Microarray Shared Resource Core Facility at University Of Colorado Denver Cancer Center. Libraries were sequenced to an average depth of 42 million reads. Reads were aligned to the human genome (hg38) using STAR Aligner (v2.7.3a) (Dobin et al., 2013), and gene counts were computed from STAR using quant mode (Dobin et al., 2013). Differential expression was performed in R using Deseq2 (Love, Huber, & Anders, 2014).

We compared our human RNAseq data to previously published datasets of KMT2D and KDM6A mutants from mice and zebrafish (Ang et al., 2016; Fahrner et al., 2019; Itoh et al., 2019; Lei & Jiao, 2018; Serrano et al., 2019; Shpargel et al., 2017). RNAseq datasets were downloaded from the NCBI Gene Expression Omnibus. Datasets included GSE129365 (Kmt2d mouse chondrocytes, (Fahrner et al., 2019)), GSE103849 (Kdm6a mouse neural crest cells, (Shpargel et al., 2017), GSE121703 and GSE128615 (mouse Cd4 cells, (Itoh et al., 2019)), GSE110392 (mouse neural stem cells, (Lei & Jiao, 2018)). In addition an RNAseq dataset of whole embryo zebrafish Kmt2d mutants was provided by Serrano et al. (Serrano et al., 2019). Where necessary, downloaded RNAseq datasets were re-analyzed using DESeq2 for differential expression.

To identify potential core genes regulated by Kmt2d and Kdm6a we found the intersection set of genes differentially expressed at a nominal p-value of 0.05 in each dataset for all Kmt2d and Kdm6a RNAseq datasets respectively. To do this, we used BioMart as implemented in R to convert non-human gene symbols into orthologous human ensembl identifiers, while allowing multiple mappings to account for gene duplications. Intersection sets were computed from these human ensemble identifiers.

### Statistical Analysis

Data shown are the means ± SEM from the number of samples or experiments indicated in the figure legends. All assays were repeated at least three times with independent samples. *P* values were determined with Student’s *t* tests using GraphPad Prism.

## Results

### Comparative analyses of the transcriptome of KS patients reveal common targets of *KMT2D* and *KDM6A*

To translate our results into the context of human birth defects, we investigated the transcriptome of 2 human Kabuki Syndrome patients carrying mutations in *KMT2D* and *KDM6A*, by RNA sequencing of their lymphoblastoid cells (LBCs). The patient with *KMT2D* mutation has a transition c.4288T>C (p.C1430R) (Figure 1A) and the patient with *KDM6A* mutation has a microdeletion in chromosome X resulting in a deletion of *KDM6A* (Figure 1A). Both mutations are predicted to result in a loss of function mutation. As controls, we used LBCs from age and ethnicity matched individuals. To ascertain differential gene expression, generated reads from RNA-seq were mapped to the human genome using STAR aligner (Dobin et al., 2013) and differential expression was performed in R with DESeq2. Differentially expressed genes (nominal p value <= 0.05) from *KMT2D* and KDM6A showed 1995 and 1917 differentially expressed genes as compared to control LBCs. Gene Ontology analysis suggests that many of these genes regulate pathways as presented in Table, consistent with both source and timing of cells that were sequenced.

We asked whether gene expression changes observed in our human RNA-seq datasets of KMT2D and KDM6A mutations are similar to those observed in datasets from the experimental model systems mouse and zebrafish. To do this, we compared RNA sequencing datasets for *Kmt2d*-mutant mice and zebrafish generated by other researchers (see methods) to our human dataset. For mice, the RNA sequencing dataset is derived from embryonic hearts (Ang et al., 2016), and the zebrafish data are derived from whole embryos (Serrano et al., 2019). Likewise, our *KDM6A* human RNA seq datasets were compared to three distinct RNA seq datasets of mice that are mutant for *Kdm6a*. The mice datasets are derived from NCCs (Shpargel et al., 2017), neural stem cells (Lei & Jiao, 2018), and CD4+ cells (Itoh et al., 2019). For *KDM6A*, the comparison is only between humans and mice, since zebrafish RNA sequencing datasets for *kdm6a* mutation are not available.

For each, we asked what genes are differentially expressed in all three datasets. A comparison between KMT2D datasets from 3 species revealed 76 orthologous genes differentially expressed in all three datasets (Figure 1B). Heat map analysis shows that 30 of the 76 common targets are either upregulated or downregulated in a similar pattern across the 3 species, while 46 do not share a similarity in expression patterns among the 3 species (Figure 1C). The dissimilarity is likely due to differences in cell type and age. For *KDM6A*, 7 common targets were found between human and the 3 mice datasets (Figure 1A). The variation in the number of common targets between *KMT2D* and *KDM6A* datasets could partly be due to the differences in the roles and targets of *KMT2D* and *KDM6A*, as well as functional redundancy of other family members across species (Akerberg, Henner, Stewart, & Stankunas, 2017; Crump & Milne, 2019; Fellous, Earley, & Silvestre, 2019; Nottke, Colaiacovo, & Shi, 2009).

Despite the dissimilarity among datasets, genes identified in these analyses may represent a set of genes commonly regulated by KMT2D and KDM6A across tissue types and taxa. We identified several genes based on known roles in regulating development with special focus on genes that affect development of craniofacial, cardiac, or NCC development including *KMT2D* datasets are *SOX12*, *SLC22A23*, *FOXP4*, *NR2F6*, and *SIX3*. The genes for *KDM6A* datasets are *ID3*, *PRDM1*, and *ANXA4*.

Comparison of the *KMT2D* and *KDM6A* datasets between human patients generated by us, with those of mice and zebrafish generated by other groups, show variations in the expression patterns of the some genes that are common across different species and cell types. The observation indicates there may be a tight spatial and temporal orchestration of the roles of *KMT2D* and *KDM6A* in gene regulation.

## Discussion

In animal models, mutations or knockdowns in Kabuki Syndrome genes show similar NCC related phenotypes. *kmt2d* mutations in zebrafish cause defects in cardiovascular development, angiogenesis and aortic arch formation, as well as cardiac hypoplasticity (Serrano et al., 2019) while Kmt2d associates with genes in cardiomyocytes, and myocardial deletion of *Kmt2d* reduces cardiac gene expression during murine heart development (Ang et al., 2016). In *Xenopus*, morpholino-based knockdown of *kmt2d* causes cardiac hypoplasticity, reduced expression of early cardiac developmental genes *tbx20*, *isl1*, *nkx2.5* (Schwenty-Lara, Nurnberger, et al., 2019), and impaired formation and migration of NCCs by reducing the expression of early NCC-specific markers like *pax3*, *foxd3*, *twist*, and *slug* (Schwenty-Lara, Nehl, et al., 2019). Interestingly we did not observe these same defects in single zebrafish Kabuki Syndrome mutants. It is likely that genetic compensation between closely related paralogs are able to compensate for the loss of function of a single gene. Future studies will test this by the creation of double and triple KS mutant zebrafish.

Less is known about *kdm6a* and its zebrafish paralog *kdm6al*. In mouse embryonic stem cells, *KDM6A* or *Utx* promotes the differentiation (ESCs) into a cardiac lineage, and serves as a co-activator of core cardiac transcription factors like *Srf, Nkx2.5*, *Tbx5*, and *Gata4* (Lee et al., 2012). *KDM6A* or *Utx* associates with cardiac-specific genes like *Anf* and *Baf60c*, to help recruit Brg1, an ATP-dependent chromatin remodeler that activates transcription of core cardiac transcription factors (Hang et al., 2010; Lee et al., 2012; Lickert et al., 2004). A neural crest-specific conditional deletion of *KDM6A* or *Utx* allele with a *Wnt1-Cre* transgene causes CHDs like patent ductus arteriosus and craniofacial defects including frontonasal hypoplasia, facial depression, and eyelid tears (Shpargel et al., 2017). In a percentage of the embryos a cleft palate is observed. Interestingly the phenotype is more severe in females likely due to the remaining copy of UTY on the Y chromosome. Because UTY has a non-functional methyltransferase, it is also possible that the ability of UTY to rescue UTX in males is methyltransferase independent (Shpargel et al., 2017). Consistent with the mouse data, zebrafish kdm6a and kdm6al mutants have minor reductions in gene expression, craniofacial cartilage and cardiac morphology as well as a reduction in histone modifications. Whether this function is methyltransferase dependent in zebrafish remains to be determined.

We present a comparison between the transcriptome of zebrafish, mice, and humans with mutations in *KMT2D,* and a comparison between transcriptomes of humans and three distinct cell populations of mice with mutations in *KDM6A*. Although our analyses involves transcriptomic datasets from different species, cell types, mutations, and developmental stages, we identify some common targets of KMT2D and KDM6A that transcend these biological differences. A subset of genes involved in cardiac development and physiology, vasculature, and overall development, appeared as common targets across all datasets for each mutation. Specifically, our analysis revealed 76 genes as common targets of *KMT2D* upon comparing our human datasets with the zebrafish and mouse datasets. 30 out of 76 genes show a similar expression pattern across the 3 species, while 46 genes do not. The dissimilar expression patterns of the 46 genes can partly be explained by the fact that the transcriptomes of the 3 species are derived from different cell populations derived during different stages. Hence, epigenetic regulation by KMT2D is tightly orchestrated spatially and temporally. On the other hand, a similar expression pattern in 3 species for the 30 genes by KMT2D indicates that its role is partly conserved. For *KDM6A*, we identified 7 common targets upon comparing human RNA-seq datasets with 3 mice datasets obtained from neural crest, neural stem, and CD4+ cells. Our *in vivo* analysis of *kmt2d* and *kdm6a* mutations in zebrafish corroborate the findings of previous studies investigating these two genes in cardiac development and disease (Ang et al., 2016; Lee et al., 2012; Schwenty-Lara, Nurnberger, et al., 2019; Serrano et al., 2019). The caveat with this type of transcriptomic analyses comparison is that the RNA-seq datasets are obtained from different sources and cell types across three species of humans, mice, and zebrafish, harboring different mutations in *KMT2D* and *KDM6A*. Despite the non-uniformity, there are some shared gene regulatory networks, and thus can serve as a resource to identify core or shared biological signatures of *KMT2D* and *KDM6A*. Although, a more direct comparison between similar mutations and cell-types are warranted in future studies, this analyses identifies the common targets of *KMT2D* and *KDM6A* across different species, cell types, and mutations.

The importance of histone modifications in cardiac and craniofacial development and function has been the focus of many recent studies (Stein et al., 2011). In mice, H3K27me3 is important for postnatal cardiac homeostasis (Delgado-Olguin et al., 2012), and the Polycomb complex regulates heart development (He et al., 2012) (Lee et al., 2012; Monroe et al., 2019). In zebrafish, H3K27me3 has recently been shown to silence structural genes in proliferating cardiomyocytes during wound invasion and regeneration of injured hearts (Ben-Yair et al., 2019). ChIP-sequencing studies on primary neonatal murine cardiomyocytes revealed that H3K4me3 is enriched, while H3K27me3 is reduced, at promoters of cardiac-specific genes like *Mef2c*, *Gata4*, and *Tbx5* (Z. Liu et al., 2016). This study further showed that early re-patterning of H3K4me3 and H3K27me3 occurs during reprogramming of induced cardiomyocytes (iCMs), which are used for modeling cardiac diseases and regeneration (Z. Liu et al., 2016). Cardiac reprogramming and epigenetics have been focused in other publications (Engel & Ardehali, 2018; Ieda et al., 2010; L. Liu, I. Lei, H. Karatas, et al., 2016; Liu, Lei, & Wang, 2016; Monroe et al., 2019). In craniofacial development, we have identified Kat2a and Kat2b as being important in regulating cartilage and bone development in both zebrafish and mice.

To conclude, several features of the roles of *KMT2D* and *KDM6A* remain to be deciphered, especially other factors that cross-talk with H3K4me3/K27me3 which are linked to craniofacial defects (Shpargel et al., 2017; Vallianatos & Iwase, 2015; Wilderman, VanOudenhove, Kron, Noonan, & Cotney, 2018) and CHDs such as *KDM1A*/*LSD1*, *KDM4A*, *KDM4C*, *EZH2*(Agger et al., 2007; Ahmed & Streit, 2018; L. Chen et al., 2012; Lee et al., 2012; Nicholson et al., 2013; Rosales, Carulla, Garcia, Vargas, & Lizcano, 2016; Willaredt, Gorgas, Gardner, & Tucker, 2012; Wu et al., 2015; Yang et al., 2019; Q. J. Zhang et al., 2011). We take a small step towards informing future studies which will be using various species to study the regulation of development and disease by *KMT2D* and *KDM6A*. Taken together, a comprehensive understanding of epigenetic regulation of heart and craniofacial development and disease, in combination with large next-generation sequencing datasets from patients that are continuously being generated and animal models will contribute to the emerging field of personalized medicine to translate epigenetics-based research from the bench to the bedside.

## Acknowledgements

We acknowledge Maria de los Angeles Serrano and H. Joseph Yost for providing the complete dataset of *kmt2d* mutant zebrafish RNA sequencing as referenced in (Serrano et al., 2019). The study was supported by Summer 2018 Postdoctoral Fellowship to R. Sen from the American Heart Association (18POST33960371), funding from Dr. K.B. Artinger (R01DE024034), and Dr. T.H. Shaikh. E.L is supported by a National Institutes of Health Ruth L. Kirschstein National Research Service Award T32 Postdoctoral Fellowship (T32CA17468).

---

**Table 1:** Characteristics relevant to cardiac and overall development.

| Genes common to *KMT2D* datasets | **<u>Table 1:</u> Characteristics relevant to cardiac and overall development** |
| --- | --- |
| *SOX12* | Expressed mice outflow tract 12.5-16.5 dpc (days post coitum), *Sox4*<sup>+/-</sup> *11*<sup>+/-</sup> *12*<sup>-/-</sup> mice have common trunk or truncus arteriosus, a CHD with single outflow tract (Bhattaram et al., 2010; Hoser et al., 2008; Paul, Harvey, Wegner, & Sock, 2014), expressed broadly throughout neural plate and migrating NCCs (Uy, Simoes-Costa, Koo, Sauka-Spengler, & Bronner, 2015) |
| *SLC22A23* | Expressed in heart and integral to mouse heart development expressed in heart, integral in mouse heart development (Aberg et al., 2012; Christoforou et al., 2008; Ekizoglu, Seven, Ulutin, Guliyev, & Buyru, 2018) |
| *FOXP4* | Mutants develop two hearts having proper chamber formation, and normal trabecular and compact myocardial development, with the same heart rate but asynchronous, lethal at E12.5 (Li, Zhou, Lu, & Morrissey, 2004; Y. Zhang et al., 2010) |
| *NR2F6* | Important for normal cardiac morphogenesis, including the development of coronary vasculature, left ventricular compact zone, and heart valves PMID: (Crispino et al., 2001; Hermann-Kleiter & Baier, 2014; Huggins, Bacani, Boltax, Aikawa, & Leiden, 2001) |
| *SIX3* | Early brain development (Inbal, Kim, Shin, & Solnica-Krezel, 2007), mutation implicated in holoprosencephaly, coloboma, cleft lip/palate (Lacbawan et al., 2009) |

**Table 2:** Characteristics relevant to cardiac and overall development

| Genes common to *KDM6A* datasets | **Table 2: Characteristics relevant to cardiac and overall development** |
| --- | --- |
| *ID3* | Knockout in various combinations with *ID1/2/4* causes ventricular septation defects, outflow tract atresia, missing heart tube-forming region, decreased heart size and function (Y. Chen et al., 2004; Cunningham et al., 2017; Fraidenraich et al., 2004; Hu, Xin, Hu, Sun, & Zhao, 2019) |
| *PRDM1* | Functions in mesoderm of second heart field, where it interacts with Tbx1, during outflow tract morphogenesis in mouse embryo (Vincent et al., 2014)) |
| *ANXA4* | Expressed in cardiac myocytes and upregulated in human failing hearts (Lewin et al., 2009; Matteo & Moravec, 2000) |

| **Table**<br>Gene Affected | Sex | Status | Effect at<br>cDNA Level | Effect at<br>Protein Level | Chromosomal Location<br>(hg19) |
| --- | --- | --- | --- | --- | --- |
| *KMT2D* | ♂ | Proband | c.4288T>C | p.C1430R | chr12:47726789(hg18) |
| *KDM6A* | ♀ | Proband | Microdeletion,<br>Gene deleted | Gene deleted | chrX:43620636-46881568 |

A
| Figure 4: Table<br>Gene Affected | Sex | Status | Effect at cDNA<br>Level | Effect at Protein<br>Level | Chromosomal Location<br>(hg19) |
| --- | --- | --- | --- | --- | --- |
| KMT2D | ♂ | Proband | c.4288T>C | p.C1430R | chr12:47726789(hg18) |
| KDM6A | ♀ | Proband | Microdeletion,<br>Gene deleted | Gene deleted | chrX:43620636-46881568 |

## References

Aberg, K., Adkins, D. E., Liu, Y., McClay, J. L., Bukszar, J., Jia, P., … van den Oord, E. J. (2012). Genome-wide association study of antipsychotic-induced QTc interval prolongation. Pharmacogenomics J, 12(2), 165–172. doi:10.1038/tpj.2010.76

Accari, S. L., & Fisher, P. R. (2015). Emerging Roles of JmjC Domain-Containing Proteins. Int Rev Cell Mol Biol, 319, 165–220. doi:10.1016/bs.ircmb.2015.07.003

Adam, M. P., Banka, S., Bjornsson, H. T., Bodamer, O., Chudley, A. E., Harris, J., … Kabuki Syndrome Medical Advisory, B. (2019). Kabuki syndrome: international consensus diagnostic criteria. J Med Genet, 56(2), 89–95. doi:10.1136/jmedgenet-2018-105625

Agger, K., Cloos, P. A., Christensen, J., Pasini, D., Rose, S., Rappsilber, J., … Helin, K. (2007). UTX and JMJD3 are histone H3K27 demethylases involved in HOX gene regulation and development. Nature, 449(7163), 731–734. doi:10.1038/nature06145

Ahmed, M., & Streit, A. (2018). Lsd1 interacts with cMyb to demethylate repressive histone marks and maintain inner ear progenitor identity. Development, 145(4). doi:10.1242/dev.160325

Akerberg, A. A., Henner, A., Stewart, S., & Stankunas, K. (2017). Histone demethylases Kdm6ba and Kdm6bb redundantly promote cardiomyocyte proliferation during zebrafish heart ventricle maturation. Dev Biol, 426(1), 84–96. doi:10.1016/j.ydbio.2017.03.030

Ali, M., Hom, R. A., Blakeslee, W., Ikenouye, L., & Kutateladze, T. G. (2014). Diverse functions of PHD fingers of the MLL/KMT2 subfamily. Biochim Biophys Acta, 1843(2), 366–371. doi:10.1016/j.bbamcr.2013.11.016

Ang, S. Y., Uebersohn, A., Spencer, C. I., Huang, Y., Lee, J. E., Ge, K., & Bruneau, B. G. (2016). KMT2D regulates specific programs in heart development via histone H3 lysine 4 di-methylation. Development, 143(5), 810–821. doi:10.1242/dev.132688

Ben-Yair, R., Butty, V. L., Busby, M., Qiu, Y., Levine, S. S., Goren, A., … Burns, C. E. (2019). H3K27me3-mediated silencing of structural genes is required for zebrafish heart regeneration. Development, 146(19). doi:10.1242/dev.178632

Bhattaram, P., Penzo-Mendez, A., Sock, E., Colmenares, C., Kaneko, K. J., Vassilev, A., … Lefebvre, V. (2010). Organogenesis relies on SoxC transcription factors for the survival of neural and mesenchymal progenitors. Nat Commun, 1, 9. doi:10.1038/ncomms1008

Bruneau, B. G. (2010). Chromatin remodeling in heart development. Curr Opin Genet Dev, 20(5), 505–511. doi:10.1016/j.gde.2010.06.008

Campeau, P. M., Lu, J. T., Dawson, B. C., Fokkema, I. F., Robertson, S. P., Gibbs, R. A., & Lee, B. H. (2012). The KAT6B-related disorders genitopatellar syndrome and Ohdo/SBBYS syndrome have distinct clinical features reflecting distinct molecular mechanisms. Hum Mutat, 33(11), 1520–1525. doi:10.1002/humu.22141

Chen, L., Ma, Y., Kim, E. Y., Yu, W., Schwartz, R. J., Qian, L., & Wang, J. (2012). Conditional ablation of Ezh2 in murine hearts reveals its essential roles in endocardial cushion formation, cardiomyocyte proliferation and survival. PLoS One, 7(2), e31005. doi:10.1371/journal.pone.0031005

Chen, Y., Mironova, E., Whitaker, L. L., Edwards, L., Yost, H. J., & Ramsdell, A. F. (2004). ALK4 functions as a receptor for multiple TGF beta-related ligands to regulate left-right axis determination and mesoderm induction in Xenopus. Dev Biol, 268(2), 280–294. doi:10.1016/j.ydbio.2003.12.035

Christoforou, N., Miller, R. A., Hill, C. M., Jie, C. C., McCallion, A. S., & Gearhart, J. D. (2008). Mouse ES cell-derived cardiac precursor cells are multipotent and facilitate identification of novel cardiac genes. J Clin Invest, 118(3), 894–903. doi:10.1172/JCI33942

Cocciadiferro, D., Augello, B., De Nittis, P., Zhang, J., Mandriani, B., Malerba, N., … Merla, G. (2018). Dissecting KMT2D missense mutations in Kabuki syndrome patients. Hum Mol Genet, 27(21), 3651–3668. doi:10.1093/hmg/ddy241

Crispino, J. D., Lodish, M. B., Thurberg, B. L., Litovsky, S. H., Collins, T., Molkentin, J. D., & Orkin, S. H. (2001). Proper coronary vascular development and heart morphogenesis depend on interaction of GATA-4 with FOG cofactors. Genes Dev, 15(7), 839–844. doi:10.1101/gad.875201

Crump, N. T., & Milne, T. A. (2019). Why are so many MLL lysine methyltransferases required for normal mammalian development? Cell Mol Life Sci, 76(15), 2885–2898. doi:10.1007/s00018-019-03143-z

Cunningham, T. J., Yu, M. S., McKeithan, W. L., Spiering, S., Carrette, F., Huang, C. T., … Colas, A. R. (2017). Id genes are essential for early heart formation. Genes Dev, 31(13), 1325–1338. doi:10.1101/gad.300400.117

Delgado-Olguin, P., Huang, Y., Li, X., Christodoulou, D., Seidman, C. E., Seidman, J. G., … Bruneau, B. G. (2012). Epigenetic repression of cardiac progenitor gene expression by Ezh2 is required for postnatal cardiac homeostasis. Nat Genet, 44(3), 343–347. doi:10.1038/ng.1068

Delgado-Olguin, P., Takeuchi, J. K., & Bruneau, B. G. (2006). Chromatin modification and remodeling in heart development. ScientificWorldJournal, 6, 1851–1861. doi:10.1100/tsw.2006.315

Digilio, M. C., Gnazzo, M., Lepri, F., Dentici, M. L., Pisaneschi, E., Baban, A., … Dallapiccola, B. (2017). Congenital heart defects in molecularly proven Kabuki syndrome patients. Am J Med Genet A, 173(11), 2912–2922. doi:10.1002/ajmg.a.38417

Dobin, A., Davis, C. A., Schlesinger, F., Drenkow, J., Zaleski, C., Jha, S., … Gingeras, T. R. (2013). STAR: ultrafast universal RNA-seq aligner. Bioinformatics, 29(1), 15–21. doi:10.1093/bioinformatics/bts635

Ekizoglu, S., Seven, D., Ulutin, T., Guliyev, J., & Buyru, N. (2018). Investigation of the SLC22A23 gene in laryngeal squamous cell carcinoma. BMC Cancer, 18(1), 477. doi:10.1186/s12885-018-4381-y

Engel, J. L., & Ardehali, R. (2018). Direct Cardiac Reprogramming: Progress and Promise. Stem Cells Int, 2018, 1435746. doi:10.1155/2018/1435746

Fahrner, J. A., Lin, W. Y., Riddle, R. C., Boukas, L., DeLeon, V. B., Chopra, S., … Bjornsson, H. T. (2019). Precocious chondrocyte differentiation disrupts skeletal growth in Kabuki syndrome mice. JCI Insight, 4(20). doi:10.1172/jci.insight.129380

Faralli, H., Wang, C., Nakka, K., Benyoucef, A., Sebastian, S., Zhuang, L., … Dilworth, F. J. (2016). UTX demethylase activity is required for satellite cell-mediated muscle regeneration. J Clin Invest, 126(4), 1555–1565. doi:10.1172/JCI83239

Fellous, A., Earley, R. L., & Silvestre, F. (2019). The Kdm/Kmt gene families in the self-fertilizing mangrove rivulus fish, Kryptolebias marmoratus, suggest involvement of histone methylation machinery in development and reproduction. Gene, 687, 173–187. doi:10.1016/j.gene.2018.11.046

Fraidenraich, D., Stillwell, E., Romero, E., Wilkes, D., Manova, K., Basson, C. T., & Benezra, R. (2004). Rescue of cardiac defects in id knockout embryos by injection of embryonic stem cells. Science, 306(5694), 247–252. doi:10.1126/science.1102612

Gagnon, J. A., Valen, E., Thyme, S. B., Huang, P., Akhmetova, L., Pauli, A., … Schier, A. F. (2014). Efficient mutagenesis by Cas9 protein-mediated oligonucleotide insertion and large-scale assessment of single-guide RNAs. PLoS One, 9(5), e98186. doi:10.1371/journal.pone.0098186

Gazova, I., Lengeling, A., & Summers, K. M. (2019). Lysine demethylases KDM6A and UTY: The X and Y of histone demethylation. Mol Genet Metab, 127(1), 31–44. doi:10.1016/j.ymgme.2019.04.012

Hang, C. T., Yang, J., Han, P., Cheng, H. L., Shang, C., Ashley, E., … Chang, C. P. (2010). Chromatin regulation by Brg1 underlies heart muscle development and disease. Nature, 466(7302), 62–67. doi:10.1038/nature09130

He, A., Ma, Q., Cao, J., von Gise, A., Zhou, P., Xie, H., … Pu, W. T. (2012). Polycomb repressive complex 2 regulates normal development of the mouse heart. Circ Res, 110(3), 406–415. doi:10.1161/CIRCRESAHA.111.252205

Hermann-Kleiter, N., & Baier, G. (2014). Orphan nuclear receptor NR2F6 acts as an essential gatekeeper of Th17 CD4+ T cell effector functions. Cell Commun Signal, 12, 38. doi:10.1186/1478-811X-12-38

Hong, S., Cho, Y. W., Yu, L. R., Yu, H., Veenstra, T. D., & Ge, K. (2007). Identification of JmjC domain-containing UTX and JMJD3 as histone H3 lysine 27 demethylases. Proc Natl Acad Sci U S A, 104(47), 18439–18444. doi:10.1073/pnas.0707292104

Hoser, M., Potzner, M. R., Koch, J. M., Bosl, M. R., Wegner, M., & Sock, E. (2008). Sox12 deletion in the mouse reveals nonreciprocal redundancy with the related Sox4 and Sox11 transcription factors. Mol Cell Biol, 28(15), 4675–4687. doi:10.1128/MCB.00338-08

Hu, W., Xin, Y., Hu, J., Sun, Y., & Zhao, Y. (2019). Inhibitor of DNA binding in heart development and cardiovascular diseases. Cell Commun Signal, 17(1), 51. doi:10.1186/s12964-019-0365-z

Huggins, G. S., Bacani, C. J., Boltax, J., Aikawa, R., & Leiden, J. M. (2001). Friend of GATA 2 physically interacts with chicken ovalbumin upstream promoter-TF2 (COUP-TF2) and COUP-TF3 and represses COUP-TF2-dependent activation of the atrial natriuretic factor promoter. J Biol Chem, 276(30), 28029–28036. doi:10.1074/jbc.M103577200

Ieda, M., Fu, J. D., Delgado-Olguin, P., Vedantham, V., Hayashi, Y., Bruneau, B. G., & Srivastava, D. (2010). Direct reprogramming of fibroblasts into functional cardiomyocytes by defined factors. Cell, 142(3), 375–386. doi:10.1016/j.cell.2010.07.002

Inbal, A., Kim, S. H., Shin, J., & Solnica-Krezel, L. (2007). Six3 represses nodal activity to establish early brain asymmetry in zebrafish. Neuron, 55(3), 407–415. doi:10.1016/j.neuron.2007.06.037

Itoh, Y., Golden, L. C., Itoh, N., Matsukawa, M. A., Ren, E., Tse, V., … Voskuhl, R. R. (2019). The X-linked histone demethylase Kdm6a in CD4+ T lymphocytes modulates autoimmunity. J Clin Invest, 130, 3852–3863. doi:10.1172/JCI126250

Kimmel, C. B., Ballard, W. W., Kimmel, S. R., Ullmann, B., & Schilling, T. F. (1995). Stages of embryonic development of the zebrafish. Dev Dyn, 203(3), 253–310. Retrieved from http://www.ncbi.nlm.nih.gov/entrez/query.fcgi?cmd=Retrieve&db=PubMed&dopt=Citation&list_uids=8589427

Klose, R. J., Kallin, E. M., & Zhang, Y. (2006). JmjC-domain-containing proteins and histone demethylation. Nat Rev Genet, 7(9), 715–727. doi:10.1038/nrg1945

Koutsioumpa, M., Hatziapostolou, M., Polytarchou, C., Tolosa, E. J., Almada, L. L., Mahurkar-Joshi, S., … Iliopoulos, D. (2019). Lysine methyltransferase 2D regulates pancreatic carcinogenesis through metabolic reprogramming. Gut, 68(7), 1271–1286. doi:10.1136/gutjnl-2017-315690

Lacbawan, F., Solomon, B. D., Roessler, E., El-Jaick, K., Domene, S., Velez, J. I., … Muenke, M. (2009). Clinical spectrum of SIX3-associated mutations in holoprosencephaly: correlation between genotype, phenotype and function. J Med Genet, 46(6), 389–398. doi:10.1136/jmg.2008.063818

Lan, F., Bayliss, P. E., Rinn, J. L., Whetstine, J. R., Wang, J. K., Chen, S., … Shi, Y. (2007). A histone H3 lysine 27 demethylase regulates animal posterior development. Nature, 449(7163), 689–694. doi:10.1038/nature06192

Lee, S., Lee, J. W., & Lee, S. K. (2012). UTX, a histone H3-lysine 27 demethylase, acts as a critical switch to activate the cardiac developmental program. Dev Cell, 22(1), 25–37. doi:10.1016/j.devcel.2011.11.009

Lei, X., & Jiao, J. (2018). UTX Affects Neural Stem Cell Proliferation and Differentiation through PTEN Signaling. Stem Cell Reports, 10(4), 1193–1207. doi:10.1016/j.stemcr.2018.02.008

Lewin, G., Matus, M., Basu, A., Frebel, K., Rohsbach, S. P., Safronenko, A., … Muller, F. U. (2009). Critical role of transcription factor cyclic AMP response element modulator in beta1-adrenoceptor-mediated cardiac dysfunction. Circulation, 119(1), 79–88. doi:10.1161/CIRCULATIONAHA.108.786533

Li, S., Zhou, D., Lu, M. M., & Morrisey, E. E. (2004). Advanced cardiac morphogenesis does not require heart tube fusion. Science, 305(5690), 1619–1622. doi:10.1126/science.1098674

Lickert, H., Takeuchi, J. K., Von Both, I., Walls, J. R., McAuliffe, F., Adamson, S. L., … Bruneau, B. G. (2004). Baf60c is essential for function of BAF chromatin remodelling complexes in heart development. Nature, 432(7013), 107–112. doi:10.1038/nature03071

Liu, L., Lei, I., Karatas, H., Li, Y., Wang, L., Gnatovskiy, L., … Wang, Z. (2016). Targeting Mll1 H3K4 methyltransferase activity to guide cardiac lineage specific reprogramming of fibroblasts. Cell Discov, 2, 16036. doi:10.1038/celldisc.2016.36

Liu, L., Lei, I., & Wang, Z. (2016). Improving cardiac reprogramming for heart regeneration. Curr Opin Organ Transplant, 21(6), 588–594. doi:10.1097/MOT.0000000000000363

Liu, Z., Chen, O., Zheng, M., Wang, L., Zhou, Y., Yin, C., … Qian, L. (2016). Re-patterning of H3K27me3, H3K4me3 and DNA methylation during fibroblast conversion into induced cardiomyocytes. Stem Cell Res, 16(2), 507–518. doi:10.1016/j.scr.2016.02.037

Love, M. I., Huber, W., & Anders, S. (2014). Moderated estimation of fold change and dispersion for RNA-seq data with DESeq2. Genome Biol, 15(12), 550. doi:10.1186/s13059-014-0550-8

Luperchio, T. R., Applegate, C. D., Bodamer, O., & Bjornsson, H. T. (2019). Haploinsufficiency of KMT2D is sufficient to cause Kabuki syndrome and is compatible with life. Mol Genet Genomic Med, e1072. doi:10.1002/mgg3.1072

Matteo, R. G., & Moravec, C. S. (2000). Immunolocalization of annexins IV, V and VI in the failing and non-failing human heart. Cardiovasc Res, 45(4), 961–970. doi:10.1016/s0008-6363(99)00409-5

Monroe, T. O., Hill, M. C., Morikawa, Y., Leach, J. P., Heallen, T., Cao, S., … Martin, J. F. (2019). YAP Partially Reprograms Chromatin Accessibility to Directly Induce Adult Cardiogenesis In Vivo. Dev Cell, 48(6), 765–779 e767. doi:10.1016/j.devcel.2019.01.017

Nicholson, T. B., Singh, A. K., Su, H., Hevi, S., Wang, J., Bajko, J., … Chen, T. (2013). A hypomorphic lsd1 allele results in heart development defects in mice. PLoS One, 8(4), e60913. doi:10.1371/journal.pone.0060913

Nottke, A., Colaiacovo, M. P., & Shi, Y. (2009). Developmental roles of the histone lysine demethylases. Development, 136(6), 879–889. doi:10.1242/dev.020966

Paul, M. H., Harvey, R. P., Wegner, M., & Sock, E. (2014). Cardiac outflow tract development relies on the complex function of Sox4 and Sox11 in multiple cell types. Cell Mol Life Sci, 71(15), 2931–2945. doi:10.1007/s00018-013-1523-x

Rosales, W., Carulla, J., Garcia, J., Vargas, D., & Lizcano, F. (2016). Role of Histone Demethylases in Cardiomyocytes Induced to Hypertrophy. Biomed Res Int, 2016, 2634976. doi:10.1155/2016/2634976

Schwenty-Lara, J., Nehl, D., & Borchers, A. (2019). The histone methyltransferase KMT2D, mutated in Kabuki syndrome patients, is required for neural crest cell formation and migration. Hum Mol Genet. doi:10.1093/hmg/ddz284

Schwenty-Lara, J., Nurnberger, A., & Borchers, A. (2019). Loss of function of Kmt2d, a gene mutated in Kabuki syndrome, affects heart development in Xenopus laevis. Dev Dyn, 248(6), 465–476. doi:10.1002/dvdy.39

Serra-Juhe, C., Cusco, I., Homs, A., Flores, R., Toran, N., & Perez-Jurado, L. A. (2015). DNA methylation abnormalities in congenital heart disease. Epigenetics, 10(2), 167–177. doi:10.1080/15592294.2014.998536

Serrano, M. L. A., Demarest, B. L., Tone-Pah-Hote, T., Tristani-Firouzi, M., & Yost, H. J. (2019). Inhibition of Notch signaling rescues cardiovascular development in Kabuki Syndrome. PLoS Biol, 17(9), e3000087. doi:10.1371/journal.pbio.3000087

Shangguan, H., Su, C., Ouyang, Q., Cao, B., Wang, J., Gong, C., & Chen, R. (2019). Kabuki syndrome: novel pathogenic variants, new phenotypes and review of literature. Orphanet J Rare Dis, 14(1), 255. doi:10.1186/s13023-019-1219-x

Shi, Y., & Whetstine, J. R. (2007). Dynamic regulation of histone lysine methylation by demethylases. Mol Cell, 25(1), 1–14. doi:10.1016/j.molcel.2006.12.010

Shpargel, K. B., Mangini, C. L., Xie, G., Ge, K., & Magnuson, T. (2020). The KMT2D Kabuki syndrome histone methylase controls neural crest cell differentiation and facial morphology. Development, 147(21). doi:10.1242/dev.187997

Shpargel, K. B., Starmer, J., Wang, C., Ge, K., & Magnuson, T. (2017). UTX-guided neural crest function underlies craniofacial features of Kabuki syndrome. Proc Natl Acad Sci U S A, 114(43), E9046–E9055. doi:10.1073/pnas.1705011114

Shpargel, K. B., Starmer, J., Yee, D., Pohlers, M., & Magnuson, T. (2014). KDM6 demethylase independent loss of histone H3 lysine 27 trimethylation during early embryonic development. PLoS Genet, 10(8), e1004507. doi:10.1371/journal.pgen.1004507

Stein, A. B., Jones, T. A., Herron, T. J., Patel, S. R., Day, S. M., Noujaim, S. F., … Dressler, G. R. (2011). Loss of H3K4 methylation destabilizes gene expression patterns and physiological functions in adult murine cardiomyocytes. J Clin Invest, 121(7), 2641–2650. doi:10.1172/JCI44641

Tekendo-Ngongang, C., Kruszka, P., Martinez, A. F., & Muenke, M. (2019). Novel heterozygous variants in KMT2D associated with holoprosencephaly. Clin Genet, 96(3), 266–270. doi:10.1111/cge.13598

Toni, L., Hailu, F., & Sucharov, C. C. (2020). Dysregulated micro-RNAs and long noncoding RNAs in cardiac development and pediatric heart failure. Am J Physiol Heart Circ Physiol, 318(5), H1308–H1315. doi:10.1152/ajpheart.00511.2019

Uy, B. R., Simoes-Costa, M., Koo, D. E., Sauka-Spengler, T., & Bronner, M. E. (2015). Evolutionarily conserved role for SoxC genes in neural crest specification and neuronal differentiation. Dev Biol, 397(2), 282–292. doi:10.1016/j.ydbio.2014.09.022

Vallianatos, C. N., & Iwase, S. (2015). Disrupted intricacy of histone H3K4 methylation in neurodevelopmental disorders. Epigenomics, 7(3), 503–519. doi:10.2217/epi.15.1

Vincent, S. D., Mayeuf-Louchart, A., Watanabe, Y., Brzezinski, J. A. t., Miyagawa-Tomita, S., Kelly, R. G., & Buckingham, M. (2014). Prdm1 functions in the mesoderm of the second heart field, where it interacts genetically with Tbx1, during outflow tract morphogenesis in the mouse embryo. Hum Mol Genet, 23(19), 5087–5101. doi:10.1093/hmg/ddu232

Vissers, L. E., van Ravenswaaij, C. M., Admiraal, R., Hurst, J. A., de Vries, B. B., Janssen, I. M., … van Kessel, A. G. (2004). Mutations in a new member of the chromodomain gene family cause CHARGE syndrome. Nat Genet, 36(9), 955–957. doi:10.1038/ng1407

Walport, L. J., Hopkinson, R. J., Vollmar, M., Madden, S. K., Gileadi, C., Oppermann, U., … Johansson, C. (2014). Human UTY(KDM6C) is a male-specific N-methyl lysyl demethylase. J Biol Chem, 289(26), 18302–18313. doi:10.1074/jbc.M114.555052

Westerfield, M. (2000). The Zebrafish Book: A Guide for the Laboratory Use of Zebrafish (Danio Rerio): University of Oregon Press.

Wilderman, A., VanOudenhove, J., Kron, J., Noonan, J. P., & Cotney, J. (2018). High-Resolution Epigenomic Atlas of Human Embryonic Craniofacial Development. Cell Rep, 23(5), 1581–1597. doi:10.1016/j.celrep.2018.03.129

Willaredt, M. A., Gorgas, K., Gardner, H. A., & Tucker, K. L. (2012). Multiple essential roles for primary cilia in heart development. Cilia, 1(1), 23. doi:10.1186/2046-2530-1-23

Wu, L., Wary, K. K., Revskoy, S., Gao, X., Tsang, K., Komarova, Y. A., … Malik, A. B. (2015). Histone Demethylases KDM4A and KDM4C Regulate Differentiation of Embryonic Stem Cells to Endothelial Cells. Stem Cell Reports, 5(1), 10–21. doi:10.1016/j.stemcr.2015.05.016

Yang, Y., Hao, H., Wu, X., Guo, S., Liu, Y., Ran, J., … Zhou, J. (2019). Mixed-lineage leukemia protein 2 suppresses ciliary assembly by the modulation of actin dynamics and vesicle transport. Cell Discov, 5, 33. doi:10.1038/s41421-019-0100-3

Yap, C. S., Jamuar, S. S., Lai, A. H. M., Tan, E. S., Ng, I., Ting, T. W., & Tan, E. C. (2020). Identification of KMT2D and KDM6A variants by targeted sequencing from patients with Kabuki syndrome and other congenital disorders. Gene, 731, 144360. doi:10.1016/j.gene.2020.144360

Zaidi, S., Choi, M., Wakimoto, H., Ma, L., Jiang, J., Overton, J. D., … Lifton, R. P. (2013). De novo mutations in histone-modifying genes in congenital heart disease. Nature, 498(7453), 220–223. doi:10.1038/nature12141

Zhang, Q. J., Chen, H. Z., Wang, L., Liu, D. P., Hill, J. A., & Liu, Z. P. (2011). The histone trimethyllysine demethylase JMJD2A promotes cardiac hypertrophy in response to hypertrophic stimuli in mice. J Clin Invest, 121(6), 2447–2456. doi:10.1172/JCI46277

Zhang, Q. J., & Liu, Z. P. (2015). Histone methylations in heart development, congenital and adult heart diseases. Epigenomics, 7(2), 321–330. doi:10.2217/epi.14.60

Zhang, Y., Li, S., Yuan, L., Tian, Y., Weidenfeld, J., Yang, J., … Morrisey, E. E. (2010). Foxp1 coordinates cardiomyocyte proliferation through both cell-autonomous and nonautonomous mechanisms. Genes Dev, 24(16), 1746–1757. doi:10.1101/gad.1929210

